# Deficiency of the membrane androgen receptor ZIP9 alters brain zinc distribution, reproductive endocrinology, and female fertility

**DOI:** 10.64898/2026.05.05.722169

**Authors:** Rui Wang, Rhiannon E. Boseley, Kalotina Geraki, Alexander P. Morrell, Alex Griffiths, Aubrey Converse, Peter Thomas, Kim C. Jonas, Robert Hindges, Christer Hogstrand

**Affiliations:** Department of Women & Children’s Health, King’s College London, London, UK; Diamond Light Source, Harwell Science and Innovation Campus, Didcot, UK; London Metallomics Facility, King’s College London, London, UK; Marine Science Institute, University of Texas at Austin, Austin, TX, USA; Centre for Developmental Neurobiology, King’s College London, London, UK; Institute of Pharmaceutical Sciences, King’s College London, London, UK

## Abstract

Zinc is an essential trace element involved in numerous biological processes, including cellular signalling, development, and reproduction. Zinc homeostasis is regulated by zinc transporters, yet the physiological roles of many transporters remain poorly understood *in vivo*. Here, we investigated the function of the zinc transporter ZIP9 (SLC39A9) using a zebrafish (*Danio rerio*) knockout model. Elemental imaging using laser ablation inductively coupled plasma mass spectrometry (LA-ICP-MS) revealed altered zinc distribution in *zip9*-deficient larvae. Synchrotron-based X-ray fluorescence (XRF) imaging further showed reduced zinc levels in the brain region of mutant zebrafish. Consistent with these observations, loss of *zip9* was associated with altered expression of key neuroendocrine genes within the hypothalamic-pituitary-gonadal (HPG) axis. Zip9 mutant females exhibited disrupted ovarian follicle development, reduced spawning rates, and decreased egg production. In addition, embryos derived from *zip9* mutant parents displayed reduced size, impaired early development, and decreased survival. Together, these findings identify ZIP9 as a regulator of zinc distribution *in vivo* and suggest that ZIP9-mediated zinc signalling contributes to reproductive regulation in zebrafish.

---

Zinc is an essential trace element that plays fundamental roles in numerous biological processes, including enzyme catalysis, protein structure stabilisation, gene expression, and cellular signalling^**1-3**^. In vertebrates, zinc is required for normal growth, development, and reproductive function, and disturbances in zinc homeostasis have been associated with a wide range of physiological and pathological conditions^**4-6**^. Beyond its structural and catalytic functions, zinc can also act as a signalling ion that dynamically regulates cellular processes such as cell proliferation and migration^**7-10**^. Given the importance of these processes, intracellular zinc levels and spatial distribution must be tightly regulated to maintain cellular and organismal homeostasis.

Cellular zinc homeostasis is primarily controlled by two families of zinc transporters: the ZIP (Zrt- and Irt-like proteins; SLC39) family and the ZnT (SLC30) family^**11-13**^. Members of the ZIP family generally increase cytosolic zinc levels by transporting zinc into the cytoplasm from either the extracellular space or intracellular compartments, whereas ZnT transporters typically reduce cytosolic zinc levels by mediating zinc efflux or sequestration into organelles^**14,15**^. Through the coordinated activity of these transporters, cells maintain appropriate zinc concentrations while allowing zinc to participate in diverse physiological functions^**16,17**^. Although the molecular properties of several ZIP transporters have been characterised, the physiological roles of many members of this family remain incompletely understood, particularly at the whole-organism level.

Among the ZIP family members, ZIP9 (SLC39A9) has attracted particular attention because of its proposed roles in zinc transport and cellular signalling^**18-20**^. ZIP9 has been reported to facilitate zinc influx across cellular membranes and regulate intracellular zinc homeostasis^**19,20**^. In addition, ZIP9 has been identified as a membrane androgen receptor, capable of mediating rapid testosterone-induced Zn^2+^ influx and signalling responses, including activation of G proteins and downstream intracellular signalling cascades, resulting in apoptosis of fish ovarian follicles and human breast cancer cells^**19-21**^.

Consistent with this idea, previous studies have shown that androgen signalling through ZIP9 can regulate apoptosis across a range of cell types, suggesting a broader role for ZIP9 in controlling cell fate decisions^**20,22,23**^. In addition, ZIP9 expression has been associated with cellular processes such as cell migration in glioblastoma cells^**10**^, supporting its involvement in regulating cell behaviour beyond classical zinc homeostasis. In addition to these signalling roles observed in cultured cells, ZIP9 has also been investigated in zebrafish, where disruption of *zip9* has been associated with ovarian phenotypes^**24,25**^. However, ZIP9 is ubiquitously expressed and the broader physiological role of ZIP9 in regulating zinc homeostasis in extragonadal tissues *in vivo* remains unclear.

This question is particularly relevant in the context of neuroendocrine control and regulation. Zinc plays important roles in neuronal signalling and endocrine function, and has been shown to influence synaptic transmission, neuronal excitability, and hormone regulation in the brain^**26,27**^. In vertebrates, reproductive function is controlled by the hypothalamic-pituitary-gonadal (HPG) axis, in which hypothalamic neurons release gonadotropin releasing hormone (GnRH) to stimulate pituitary secretion of follicle stimulating hormone (FSH) and luteinising hormone (LH), ultimately regulating gonadal development and gametogenesis^**28,29**^. Upstream regulators such as kisspeptin integrate physiological cues to control GnRH activity^**30**^. Zinc has also been implicated in several reproductive processes, including oocyte maturation, fertilisation, and early embryonic development^**31-33**^. However, whether zinc transporters contribute to the regulation of neuroendocrine reproductive pathways remains largely unknown.

Zinc distribution within tissues is highly heterogeneous and cannot be fully captured by bulk measurements alone. Advances in elemental imaging techniques, such as laser ablation inductively coupled plasma mass spectrometry (LA-ICP-MS) and synchrotron-based X-ray fluorescence (XRF) imaging, now enable spatially resolved analysis of metal distribution in biological tissues^**34-36**^. These approaches provide an opportunity to investigate how disruption of zinc transporters may alter tissue-level metal distribution, particularly within neuroendocrine regions of the brain that regulate reproductive function.

The zebrafish (*Danio rerio*) is a widely used vertebrate model for studying developmental biology, genetics, and reproductive physiology^**37-39**^. Using a zebrafish genetic model, we examined the role of ZIP9 in zinc homeostasis and neuroendocrine regulation of reproduction in vivo. By combining a *zip9* knockout line with spatial elemental imaging, we characterised changes in zinc distribution during development and assessed the impact of *zip9* deficiency on HPG axis gene expression, ovarian morphology, reproductive performance, and early embryonic development. Together, these findings provide new insights into the role of ZIP9 in regulating zinc homeostasis and suggest that ZIP9-mediated zinc signalling contributes to the neuroendocrine control of reproduction in vertebrates

## Results

### Zip9 is expressed during zebrafish development and enriched in neuroendocrine tissues

To examine the temporal expression of *zip9* during zebrafish development, mRNA and protein levels were analysed in wildtype larvae. Quantitative PCR analysis showed that *zip9* mRNA was detectable throughout early development and gradually increased from 1 to 5 days post-fertilisation (dpf) (Fig. 1A). Consistent with this pattern, Western blot analysis indicated that Zip9 protein levels were relatively stable between 1 and 3 dpf but increased at 5 dpf (Fig. 1B). To assess the spatial distribution of Zip9 during larval development, whole-mount immunofluorescence staining was performed in 5 dpf larvae. Zip9 signal was detected throughout the body, with stronger signal intensity in the anterior region of the larvae (Fig. 1C). Elevated immunoreactivity was observed in regions located between the ventral sides of the eyes, corresponding approximately to the hypothalamic-pituitary region of the brain. Additional Zip9 signals were also detected in the muscle and caudal fin (Fig. 1C). To determine whether *zip9* is expressed in adult reproductive tissues, mRNA levels were analysed in adult zebrafish organs. Quantitative PCR revealed that *zip9* transcript was present in the brain, liver, ovary, and testis (Fig. 1D). Among these tissues, the ovary showed the highest zip9 expression. Comparison between male- and female-derived tissues indicated higher expression levels in female tissues.

**Figure 1.**
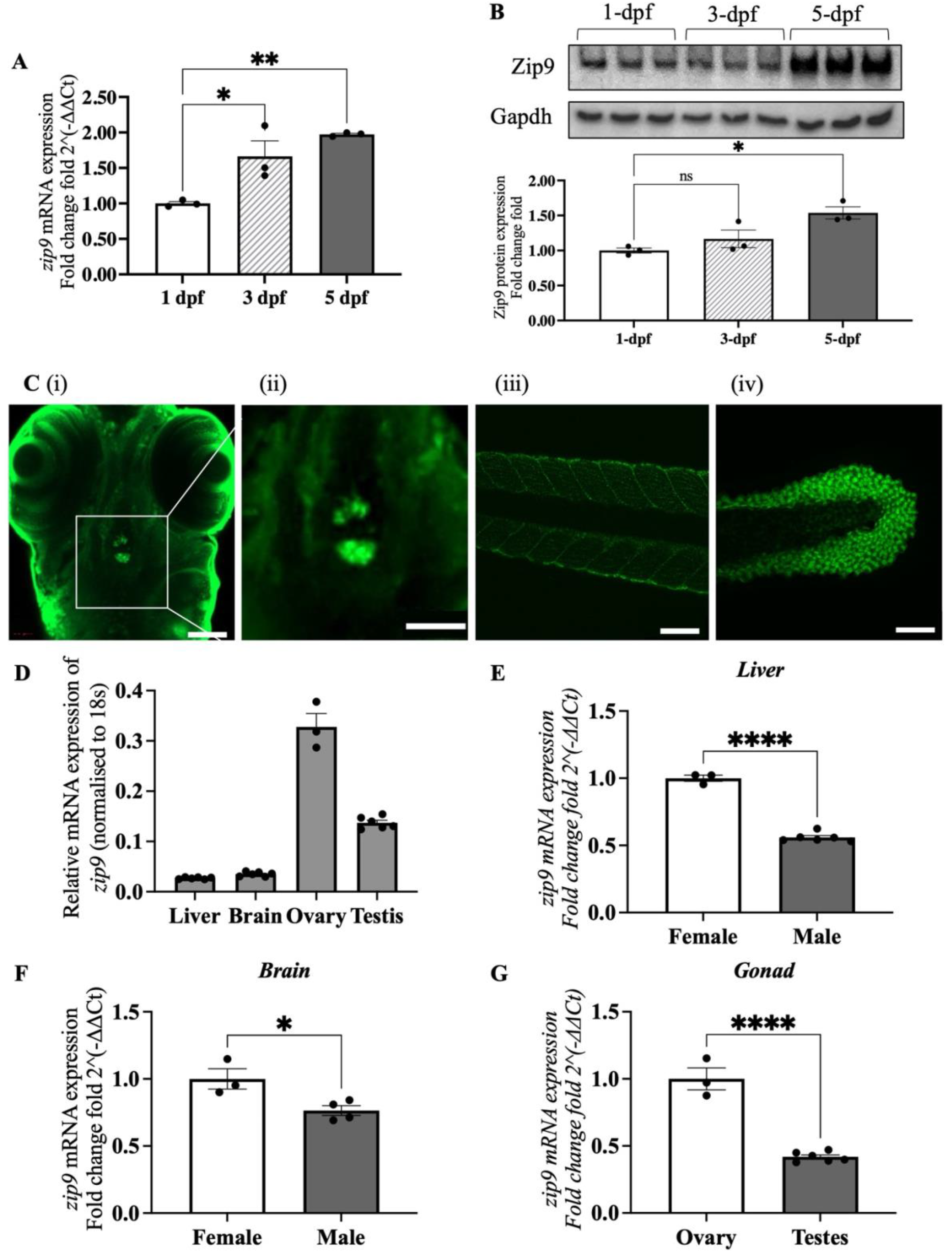
Expression of Zip9 in wildtype zebrafish. (A) Relative mRNA expression of *zip9* in wildtype zebrafish larvae at 1-, 3-, and 5-days post-fertilisation (dpf) measured by quantitative PCR, normalised to *18s*, compared to 1dpf. (B) Representative Western blot showing Zip9 and Gapdh protein levels in 1, 3, and 5 dpf larvae, quantification normalised to Gapdh, compared to 1dpf. (C) Whole-mount immunofluorescence showing Zip9 localisation in 5 dpf zebrafish larvae, including (i) head region, (ii) enlarged view of the ‘pituitary-like’ region, (iii) muscle, and (iv) caudal fin. Scale bars: 100 μm, inset 50 μm. (D) Relative *zip9* mRNA expression in adult zebrafish tissues including liver, brain, ovary, testis, and kidney, normalised to *18s*. (E-H) Sex-specific comparison of *zip9* mRNA expression in adult zebrafish tissues including liver (F), brain (G), and gonad (H), normalised to *18s*. Data are presented as mean ± SEM (n = 3). Statistical analysis was performed using one-way ANOVA (\**P* < 0.05, \*\**P* < 0.005, \*\*\**P* < 0.0001).

### Generation of a *zip9* knockout zebrafish line

To investigate the physiological role of Zip9 in zebrafish, a *zip9* knockout line was generated using the CRISPR/Cas9 genome editing system. A four-base-pair insertion was introduced into exon 1 of the *zip9* gene, resulting in a frameshift mutation that generated a premature stop codon (TAA). This mutation was predicted to truncate the Zip9 protein at amino acid position VAL20 and remove the majority of the coding sequence, including most of the predicted transmembrane domains. Genotyping by Sanger sequencing confirmed the presence of the mutation and allowed identification of wildtype (*zip9*^*+/+*^), heterozygous (*zip9*^*+/−*^), and homozygous mutant (*zip9*^*−/−*^) zebrafish. Representative sequencing chromatograms and sequence alignments confirming the *zip9* mutation are shown in Supplementary Fig. S1. These results confirm the successful generation of a *zip9* mutant zebrafish line for subsequent functional analyses.

### Loss of *zip9* alters zinc distribution in zebrafish larvae

To determine whether *zip9* deficiency affects metal distribution during early development, laser ablation inductively coupled plasma mass spectrometry (LA-ICP-MS) was performed on 5 dpf zebrafish larvae. Because homozygous embryos from *zip9*^*−/−*^ in-crosses failed to develop beyond early stages, larvae generated from crosses between *zip9*^*−/−*^ males and wildtype females were analysed to obtain *zip9*^*+/−*^ offspring. Relative analysis revealed that zinc levels were significantly reduced in *zip9*^*+/−*^ larvae compared with wildtype controls when normalised to both phosphorus and sulphur as the proxy for cell density and proteins respectively (Fig. 2A). A moderate decrease in iron signal was also observed, whereas copper and manganese levels remained comparable between genotypes (Fig. 2B-D). These results indicate that *zip9* deficiency is associated with altered zinc homeostasis during zebrafish larval development.

**Figure 2.**
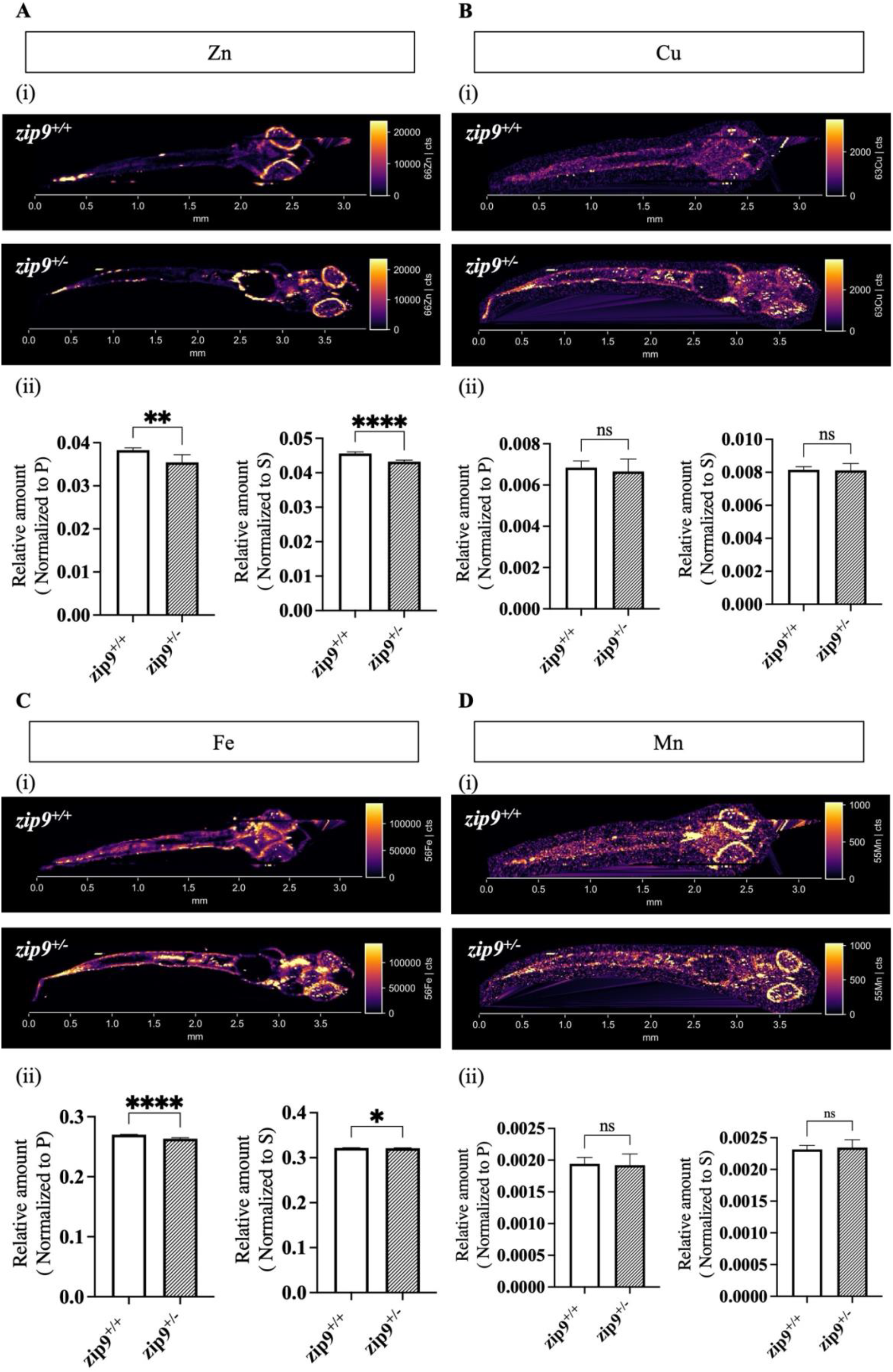
Elemental quantification in 5 dpf zebrafish larvae measured by LA-ICP-MS. Relative analysis of zinc (A), copper (B), iron (C), and manganese (D) in 5 dpf zebrafish larvae. (i) Representative LA-ICP-MS elemental maps of zebrafish larvae showing *zip9*+/+ (top) and *zip9*^*+/−*^ (bottom) samples. (ii) Relative analysis of elemental signal intensity in the whole larvae, normalised to phosphorus (left) and sulphur (right). Data are presented as mean ± SEM (n ≥ 3). Statistical analysis was performed using t test (\**P* < 0.05, \*\**P* < 0.005, \*\*\**P* < 0.0001).

In wildtype larvae, zinc signal intensity showed apparent enrichment in the anterior region of the body, including the brain region. Enlarged views suggested higher signal intensity within the central brain area (Fig. 3A). In contrast, this regional enrichment appeared less pronounced in *zip9*^*+/−*^ larvae, where the zinc signal was more diffuse (Fig. 3B). It should be noted that localised high-intensity signals observed in some regions may reflect technical artefacts associated with sample preparation or ablation, and therefore interpretation is focused on overall distribution patterns rather than isolated signal hotspots.

**Figure 3.**
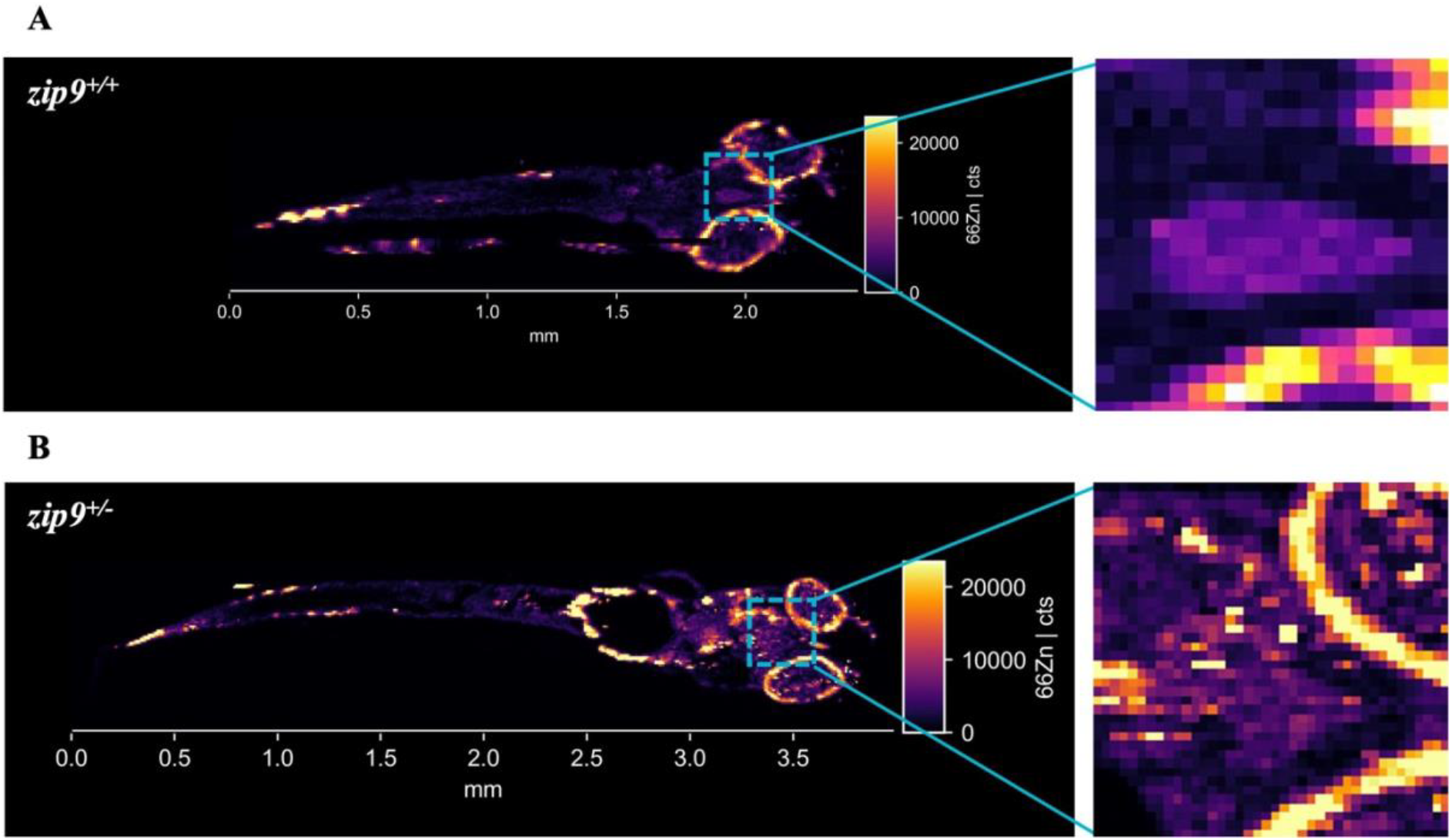
Zinc distribution in zebrafish larvae visualised by LA-ICP-MS. Representative LA-ICP-MS zinc maps of 5 dpf zebrafish larvae showing *zip9*^*+/+*^ (A) and *zip9*^*+/−*^ (B) samples. Enlarged views of the brain region are shown as insets (n ≥ 3).

### Zip9 deficiency reduces zinc levels in the zebrafish brain

To further examine the spatial distribution of zinc in the brain region following the global changes observed by LA-ICP-MS, XRF imaging was performed on heads of wildtype and *zip9*^*−/−*^ zebrafish. While LA-ICP-MS provided higher-throughput, semi-quantitative analysis of whole-body elemental distribution, XRF imaging enabled higher-resolution spatial visualisation of metal distribution within specific anatomical regions. Two-dimensional scans were acquired in two orientations (0°, dorsal view; 90°, lateral view) to visualise elemental distribution from complementary perspectives (Fig. 4). Elemental maps revealed the spatial distribution of zinc and calcium across the scanned regions of the head (Fig. 4A-B), and additional individual XRF maps used for region-of-interest (ROI) analysis are shown in Supplementary Fig. S4. Quantification of zinc intensity within selected ROIs revealed reduced zinc signal in *zip9*^*−/−*^ samples compared with wildtype controls. This reduction was observed in the hypothalamus-pituitary region, whereas the eye region, used as an anatomical reference for localisation, showed no detectable difference between genotypes (Fig. 4iii-iv). Quantitative analysis showed a significant decrease in zinc levels in the hypothalamus-pituitary region in the 90° view, while the reduction in the 0° view showed a similar trend but did not reach statistical significance (*P* = 0.0748).

**Figure 4.**
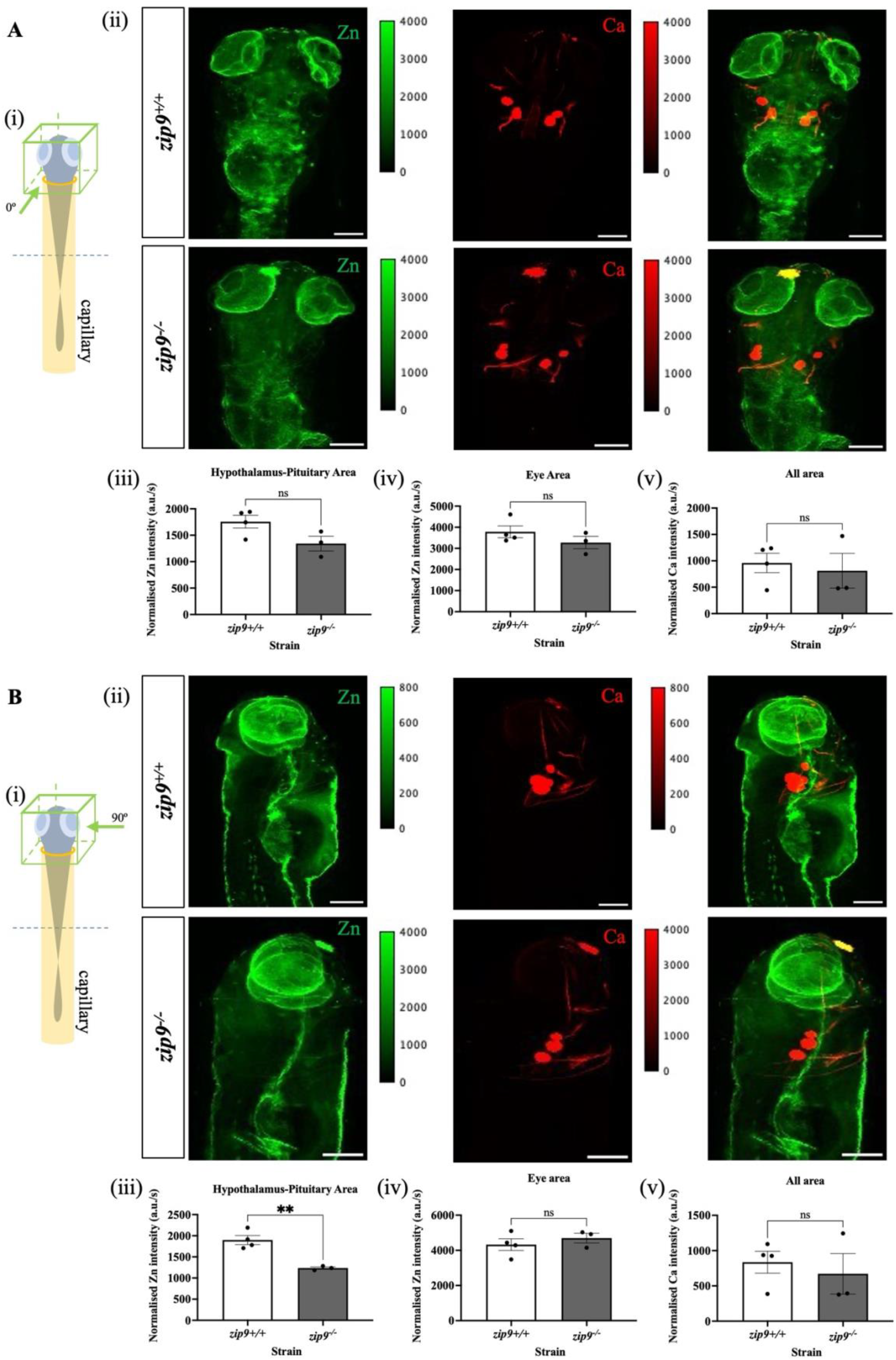
Two-dimensional X-ray fluorescence (XRF) imaging of zinc distribution in zebrafish heads. Panels A and B show scans acquired at 0° (A) and 90° (B) rotation angles. (i) Schematic illustration showing the positioning of the zebrafish head within the capillary during scanning; the 0° orientation was defined as the dorsal view position in which the anterior side of the head faced forward with both eyes symmetrically visible; the 90° orientation was defined as the lateral view position in which the anterior side of the head faced forward with both eyes overlapping. (ii) Elemental maps of zinc and calcium in *zip9*^*+/+*^ (top) and *zip9*^*−/−*^ (bottom) zebrafish heads, with merged images shown on the right. Scale bar = 0.2 mm. (iii) Quantification of zinc intensity in the hypothalamus-pituitary region. (iv) Quantification of zinc intensity in the eye region. (v) Quantification of calcium intensity across the entire scanned area. Data are presented as mean ± SEM (n ≥ 3). Statistical analysis was performed using t test (\*\**P* < 0.005).

In contrast, zinc levels in the eye region were not significantly different between genotypes in either orientation. Consistent with this regional effect, calcium intensity measured across the entire scanned area remained comparable between genotypes (Fig. 4v).

To further visualise zinc distribution within the zebrafish head, three-dimensional XRF tomography was performed by adding a rotation about the centre of the samples (Fig. 5). Three orthogonal views (front, bottom, and right-side) enabled spatial visualisation of zinc distribution within the head and brain regions. Representative three-dimensional reconstructions revealed clear differences in zinc distribution between *zip9*^*+/+*^ and *zip9*^*−/−*^ zebrafish (Fig. 5B-C). In wildtype sample, a distinct layer of elevated zinc signal was observed in the central region of the head. This high-zinc region was clearly visible in both the bottom and right-side views, where it appeared as a layer located along the central midline of the head. In contrast, this high-zinc region was absent in *zip9*^*−/−*^ sample. This location is consistent with the hypothalamus-pituitary region identified in the two-dimensional XRF analysis. Three-dimensional reconstructions of zinc distribution are provided in Supplementary Videos S1 and S2.

**Figure 5.**
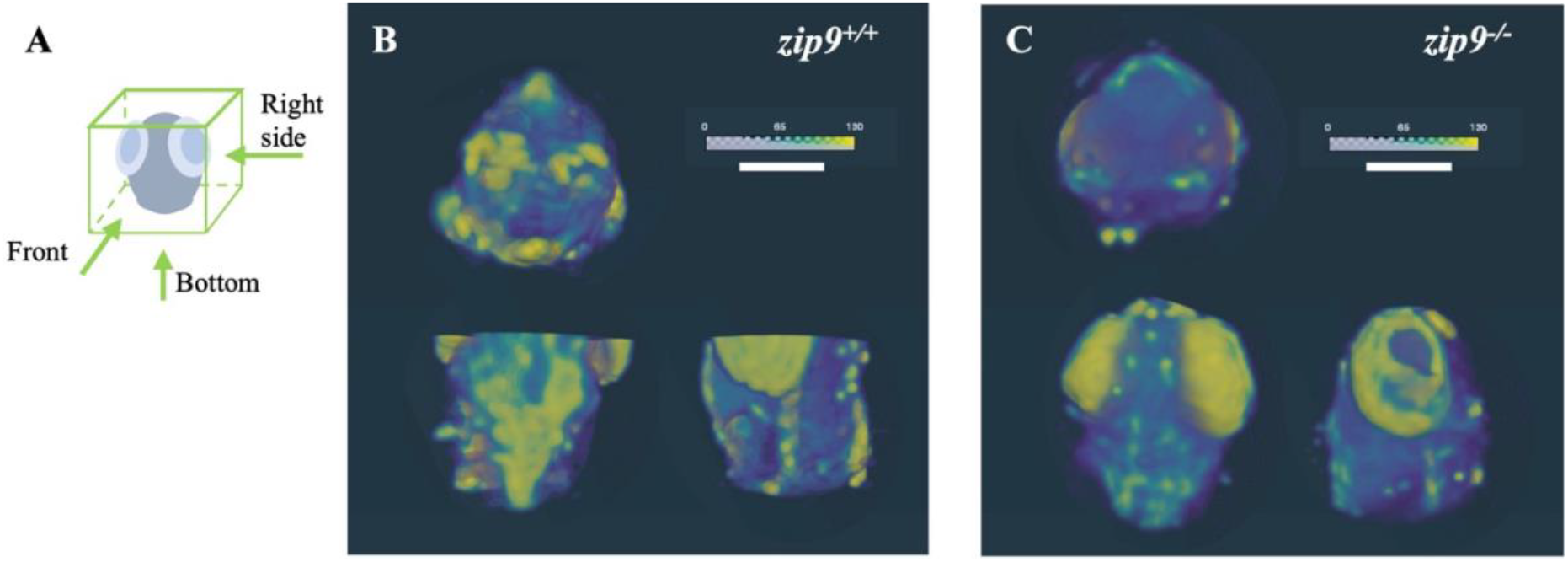
Three-dimensional X-ray fluorescence (XRF) tomography of zinc distribution in zebrafish heads. (A) Schematic illustration showing the orientation used for three-dimensional XRF scanning of the zebrafish head. (B) Three-dimensional XRF scanning showing zinc distribution in the head of a *zip9*^*+/+*^ zebrafish from three orthogonal views (front, bottom, and right-side). (C) Three-dimensional XRF scanning showing zinc distribution in the head of a *zip9*^*−/−*^ zebrafish from the same three orthogonal views. Scale bar = 0.4 mm.

### Zip9 deficiency disrupts neuroendocrine regulation of the HPG axis

Given the reduction of zinc levels in the hypothalamus-pituitary region combined with the high expression of Zip9 in the same region, the expression of key genes of the hypothalamic-pituitary-gonadal (HPG) axis was analysed in the brains-pituitary of adult zebrafish using quantitative PCR. In female zebrafish, several neuroendocrine regulatory genes showed reduced expression in *zip9*^*−/−*^ fish compared with wildtype controls. These included *kiss1* and *kiss2*, which encode neuropeptides involved in the stimulation of gonadotropin-releasing hormone (GnRH) secretion, as well as gonadotropic releasing hormone receptors 1 and 2, *gnrh2* and *gnrh3* (Fig. 6A-D). Similarly, the expression of the pituitary gonadotropin subunits *fshb* and *lhb* was also decreased in *zip9*^*−/−*^ females (Fig. 6E,F). In contrast, male *zip9*^*−/−*^ zebrafish showed an overall increase in the expression of these genes compared with wildtype males (Fig. 6A-F), indicating a sex-dependent alteration in neuroendocrine gene expression following *zip9* deficiency.

**Figure 6.**
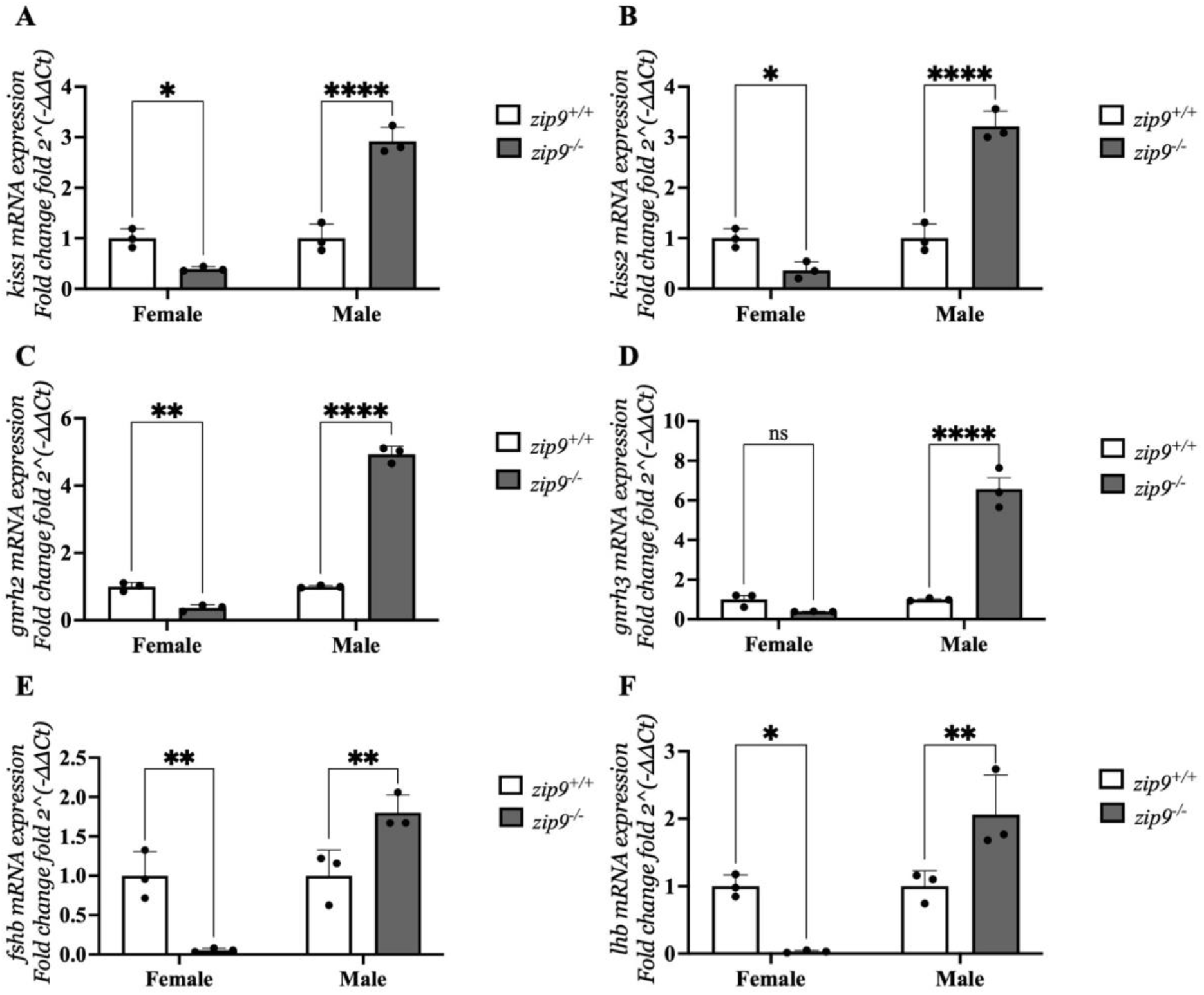
Zip9 deficiency alters neuroendocrine gene expression in the zebrafish brain-pituitary. Relative mRNA expression of key HPG axis regulatory genes in the brains of adult zebrafish measured by quantitative PCR. Expression levels of *kiss1* (A), *kiss2* (B), *gnrh2* (C), *gnrh3* (D), *fshb* (E), and *lhb* (F) are shown for wildtype (*zip9*^*+/+*^) and *zip9*^*−/−*^ fish. Data are presented separately for female and male zebrafish. Expression levels were normalised to the reference gene and are presented as mean ± SEM (n = 3). Statistical analysis was performed using two-way ANOVA(* *P* <0.05, ** *P* <0.005, **** *P* <0.0001).

### Zip9 deficiency alters gonadal and hepatic reproductive gene expression

To further examine downstream reproductive signalling, the expression of key genes involved in gonadal and hepatic reproductive regulation was analysed in adult zebrafish. In the gonads, male *zip9*^*−/−*^ zebrafish exhibited increased expression of androgen receptor (*ar*) and oestrogen receptor (*esr1*) compared with wildtype controls, same trends were observed in female *zip9*^*−/−*^ zebrafish (Fig. 7A,B). Expression of *cyp19a1*, which encodes aromatase responsible for converting androgens to oestrogens, was also elevated in *zip9*^*−/−*^ fish (Fig. 7C). Similarly, *fshr*, a receptor involved in follicle development and gametogenesis, showed increased expression in both sexes (Fig. 7D). In contrast, *lhr* expression decreasing trend in *zip9*^*−/−*^ females but significantly increased in *zip9*^*−/−*^ males (Fig. 7E). Additionally, *yap1*, a regulator of ovarian follicle development and cell proliferation, was reduced in *zip9*^*−/−*^ females, while remaining unchanged in males (Fig. 7F). To assess hepatic responses related to reproduction, expression of oestrogen receptor (*esr1*) and vitellogenin (*vtg*) was analysed in female livers. Both genes were significantly reduced in *zip9*^*−/−*^ females compared with wildtype controls (Fig. 7G,H), indicating altered hepatic regulation of yolk protein production.

**Figure 7.**
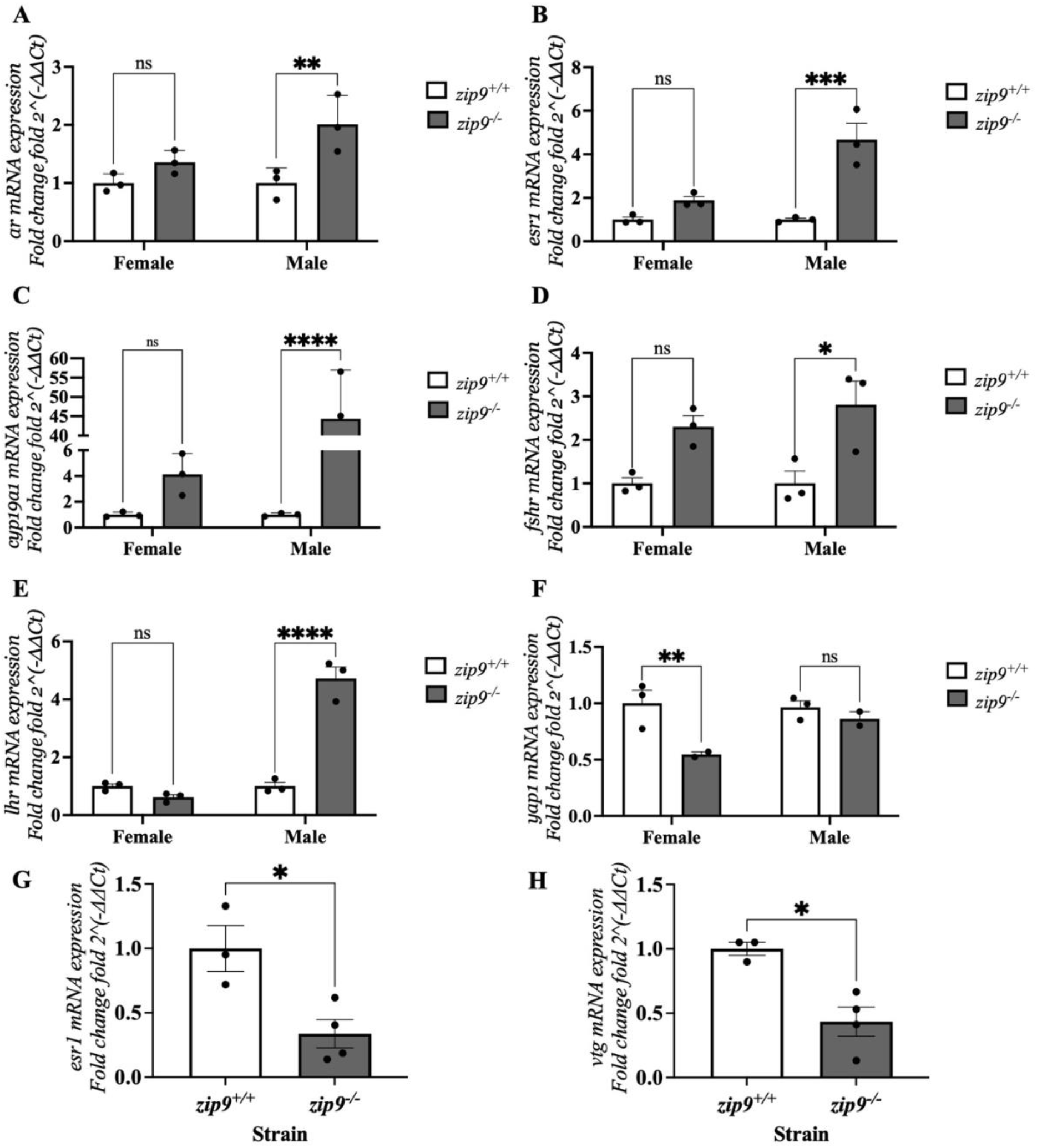
Zip9 deficiency alters gonadal and hepatic reproductive gene expression. Relative mRNA expression of reproductive regulatory genes in adult zebrafish measured by quantitative PCR. Gonadal expression levels of *ar* (A), *esr1* (B), *cyp19a1* (C), *fshr* (D), *lhr* (E), and *yap1* (F) are shown for wildtype (*zip9*^*+/+*^) and *zip9*^*−/−*^ fish, statistical analysis was performed using two-way ANOVA (* *P* <0.05, ** *P* <0.005, *** *P* <0.001, **** *P* <0.0001). Hepatic expression of *esr1* (G) and *vtg* (H) is shown in female zebrafish, statistical analysis was performed using t test (* *P* <0.05). Expression levels were normalised to the reference gene *18s* and are presented as mean ± SEM (n = 3).

### Zip9 deficiency alters gonadal morphology and follicle development

To assess whether *zip9* deficiency affects gonadal structure, histological analysis of ovaries from adult zebrafish was performed using haematoxylin and eosin staining. Wildtype ovaries were densely packed with large, mature follicles, whereas *zip9*^*−/−*^ ovaries contained a higher number of smaller follicles (Fig. 8A,B). Structural disorganisation of the somatic tissue was also observed in *zip9*^*−/−*^ ovaries, including clusters of apoptotic cells and regions of increased eosinophilic staining. Quantification of follicle number revealed a significantly higher number of follicles in *zip9*^*−/−*^ ovaries compared with wildtype controls (Fig. 8C). However, despite the increased follicle number, follicle diameters were significantly smaller in *zip9*^*−/−*^ ovaries (Fig. 8D). Consistent with this observation, follicle staging based on size distribution indicated that *zip9*^*−/−*^ ovaries contained a higher proportion of early-stage follicles, whereas wildtype ovaries showed a greater proportion of late-stage follicles (Fig. 8E). Testicular morphology was also examined in adult males (Supplementary Figure S5). Histological sections revealed that seminiferous lobules in *zip9*^*−/−*^ testes appeared smaller than those in wildtype controls, although abundant spermatozoa were present within the lobules. These observations suggest that male fertility is largely preserved in *zip9*^*−/−*^ zebrafish, consistent with the largely normal testicular morphology observed in histological analysis (Supplementary Fig. S5).

**Figure 8.**
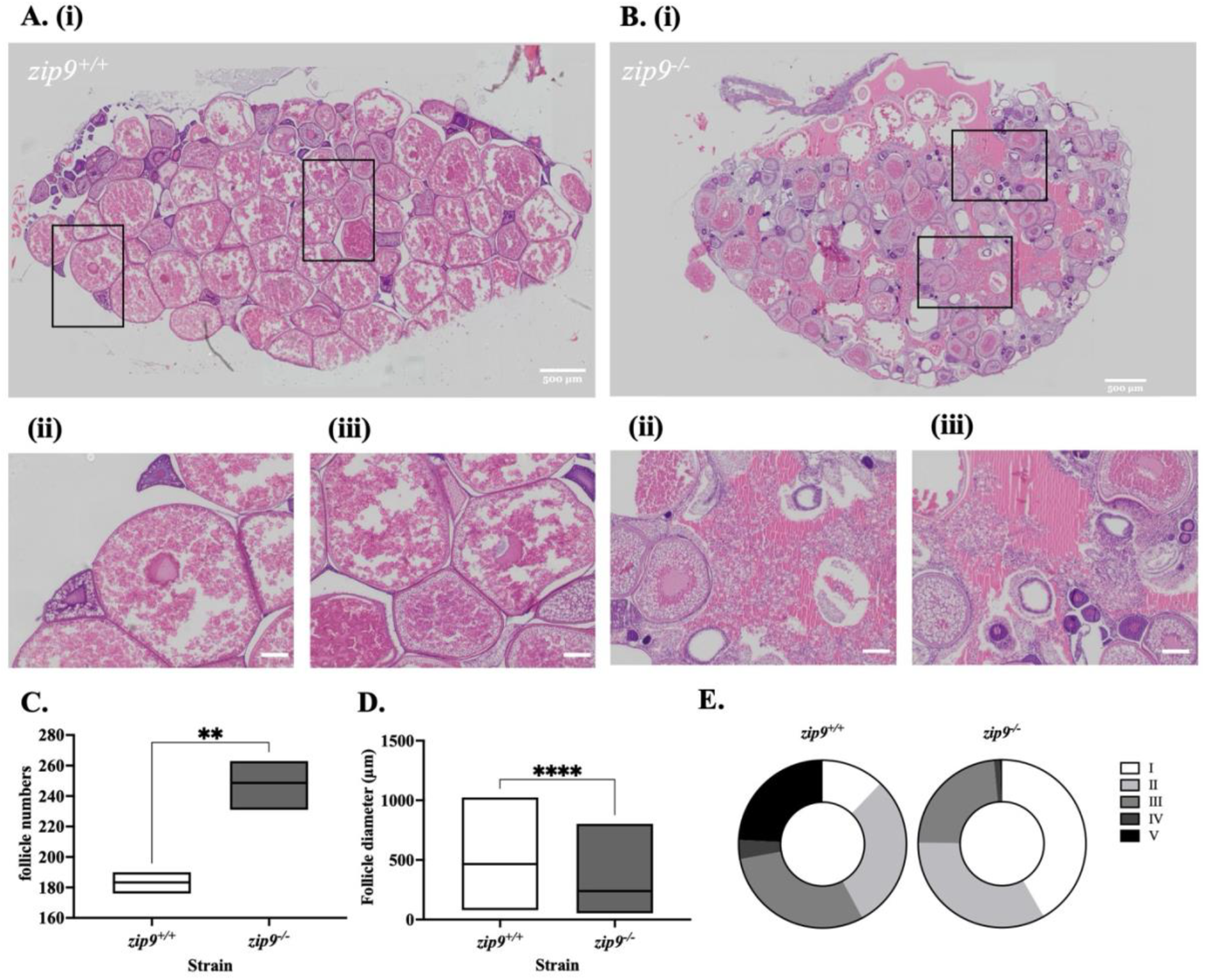
Zip9 deficiency alters ovarian morphology and follicle development. (A,B) Representative haematoxylin and eosin-stained sections of ovaries from wildtype (*zip9*^*+/+*^) (A) and *zip9*^*−/−*^ (B) adult zebrafish. (C) Quantification of ovarian follicle diameter. (D) Distribution of follicle stages based on follicle size. Follicles were classified according to diameter as stage I (7-140 µm), stage II (140-340 µm), stage III (340-690 µm), stage IV (690-730 µm), and stage V (730-750 µm). Data are presented as mean ± SEM (n = 3). Statistical analysis was performed using unpaired t-tests (\*\**P* < 0.05, \*\*\*\**P* < 0.0001).

### Zip9 deficiency reduces reproductive performance and compromises early embryonic development

To investigate whether *zip9* deficiency affects reproductive success, breeding assays were performed using wildtype (*zip9*^*+/+*^) and homozygous mutant (*zip9*^*−/−*^) zebrafish. Although *zip9*^*−/−*^ breeding pairs displayed normal courtship behaviours, including quivering and hooking, their reproductive success was significantly reduced. The spawning rate of *zip9*^*−/−*^ in-crosses was significantly lower than that of wildtype pairs, and the pairs that did spawn produced significantly fewer eggs (Fig. 9A,B). To determine whether the reproductive impairment was associated with the male or female genotype, reciprocal crosses between wildtype and *zip9*^*−/−*^ fish were performed. Crosses between *zip9*^*−/−*^ males and wildtype females resulted in spawning rates and egg numbers comparable to those observed in wildtype controls, indicating that male fertility is largely preserved in *zip9*^*−/−*^ zebrafish. In contrast, crosses between *zip9*^*−/−*^ females and wildtype males exhibited reduced spawning success and egg production similar to those observed in *zip9*^*−/−*^ in-crosses, indicating that the reproductive phenotype is primarily associated with *zip9* deficiency in females (Fig. 9A,B). Early embryonic development was subsequently examined. Wildtype embryos progressed through normal developmental stages during the first 24 hours post-fertilisation (Fig. 9C). In contrast, embryos derived from *zip9*^*−/−*^ in-crosses exhibited severe developmental impairment, with all the embryos failing to progress beyond early cleavage stages and undergoing degeneration within the first 24 hours. Measurement of chorion diameter immediately after spawning showed that embryos derived from *zip9*^*−/−*^ parents were significantly smaller than those from wildtype fish (Fig. 9D). Consistent with these observations, embryo survival at 24 hours post-fertilisation was markedly reduced in *zip9*^*−/−*^ embryos compared with wildtype controls (Fig. 9E).

**Figure 9.**
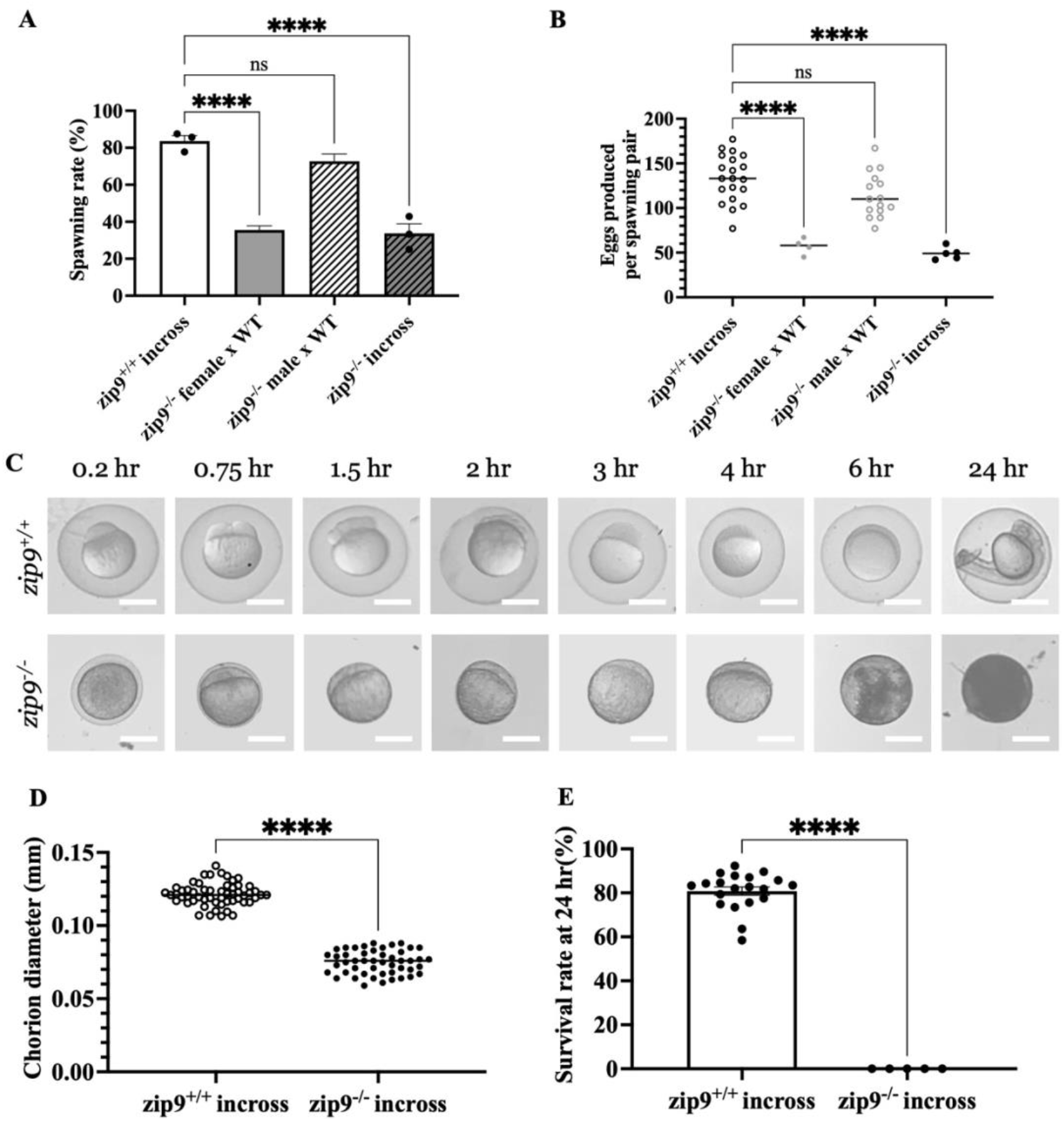
Reproductive performance and embryonic development in zip9 mutant zebrafish. (A) Spawning rate of breeding pairs across different mating combinations: wildtype in-crosses (*zip9*^*+/+*^ × *zip9*^*+/+*^), zip9 mutant in-crosses (*zip9*^*−/−*^ × *zip9*^*−/−*^), and reciprocal crosses between wildtype and *zip9*^*−/−*^ fish. (B) Number of eggs produced per spawning pair across the four mating combinations. (C) Representative images of embryonic development from wildtype and *zip9*^*−/−*^ in-crosses during the first 24 hours post-fertilisation. (D) Chorion diameter of embryos immediately after spawning. (E) Embryo survival at 24 hours post-fertilisation. Data are presented as mean ± SEM. Statistical analysis was performed using unpaired t-tests or one-way ANOVA as appropriate. n ≥ 14 independent breeding pairs.

## Discussion

In this study, we investigated the physiological function of the zinc transporter ZIP9 in zebrafish using a genetic knockout model. Loss of *zip9* resulted in altered zinc distribution during larval development, including reduced zinc levels in the brain region. This was associated with disruption of neuroendocrine gene expression within the hypothalamic-pituitary-gonadal (HPG) axis, altered reproductive gene regulation in gonadal and hepatic tissues, and defects in ovarian morphology, reproductive output and early embryonic development. Together, these findings identify Zip9 as a regulator of zinc distribution within the HPG axis that was associated with impaired neuroendocrine control of reproduction in zebrafish.

In addition to these physiological findings, the present study provides one of the first spatially resolved elemental maps of the vertebrate brain at larval stages. Using complementary LA-ICP-MS and synchrotron-based X-ray fluorescence (XRF) imaging, we identified a distinct and reproducible pattern of zinc enrichment in the central brain region. This observation is consistent with previous synchrotron XRF studies in early zebrafish embryos, suggesting that spatial organisation of zinc in the developing brain is a conserved feature^**40,41**^. Notably, zinc provided the most cohesive structural representation of the zebrafish head, highlighting its dual role as both a structural and signalling element and supporting its use as a sensitive marker for neuroanatomical organisation.

Zinc homeostasis in vertebrates is tightly regulated by ZIP (SLC39) and ZnT (SLC30) transporters^**14,42,43**^. Among these, ZIP9 is unique in that it has been proposed to function both as a zinc transporter and a membrane androgen receptor. Previous studies have shown that ZIP9 mediates rapid, non-genomic androgen signalling through G protein-dependent pathways, leading to changes in intracellular zinc levels and regulation of apoptosis in ovarian follicle cells^**19,44**^. These responses are context-dependent, with both pro- and anti-apoptotic effects observed depending on follicle stage^**45**^. *In vivo* studies further demonstrate roles for ZIP9 in ovarian homeostasis, including post-ovulatory follicle breakdown and reproductive function, as well as zinc-dependent processes such as egg activation and early embryonic development^**24,25**^. Together, these findings highlight the dual functionality of ZIP9, although the relative contribution of its zinc transport and androgen receptor activities remains unresolved.

In the present study, *zip9* deficiency resulted in reduced zinc levels in the brain, particularly in regions corresponding to the hypothalamic-pituitary area. Given the importance of zinc in neuronal signalling and endocrine regulation, disruption of ZIP9-mediated zinc transport is likely to affect neuroendocrine pathways controlling reproduction^**30**^. Consistent with this, *zip9* mutants showed altered expression of key HPG axis genes, including *kiss1, kiss2, gnrh2, gnrh3, fshb* and *lhb*, suggesting impaired neuroendocrine signalling.

In addition to central effects, *zip9* deficiency altered gene expression in reproductive tissues and was associated with impaired ovarian development, characterised by increased numbers of smaller follicles and a shift towards early-stage follicle populations. Previous studies have reported reduced fertilisation success, impaired embryonic development and delayed post-ovulatory follicle breakdown in *zip9*-deficient zebrafish^**24,25**^. While those studies focused on post-ovulatory processes, the reproductive defects observed here are consistent with a broader role for ZIP9 in female reproductive function. In contrast, male *zip9* mutants remained fertile despite modest histological differences, indicating that the phenotype is more pronounced in females, potentially reflecting the greater dependence of oocyte development on tightly regulated zinc homeostasis. However, the extent to which these effects are mediated by zinc-dependent versus androgen-dependent mechanisms remains unclear.

The reproductive defects observed in *zip9* mutants, including reduced spawning, smaller eggs and impaired embryonic development, further support a role for ZIP9 in oocyte quality and early embryogenesis. These phenotypes may reflect altered maternal provisioning associated with disrupted zinc homeostasis, which is known to play critical roles in cell cycle progression and transcriptional regulation during early development^**31-33**^.

Although this study identifies ZIP9 as a regulator of zinc distribution and reproductive physiology in zebrafish, several questions remain. In particular, the cellular mechanisms linking ZIP9-dependent zinc transport to neuroendocrine and reproductive outcomes are not yet defined. Future studies using cell-type-specific approaches and high-resolution imaging will be required to distinguish between central and local effects and to determine whether ZIP9 functions primarily through zinc-dependent signalling, androgen-dependent pathways, or an integration of both.

Together, these findings demonstrate that ZIP9 regulates zinc homeostasis in the developing brain and reproductive tissues and suggest that disruption of ZIP9-mediated zinc signalling impairs neuroendocrine control of reproduction and early embryonic development in zebrafish.

## Materials and Methods

### Zebrafish husbandry

Zebrafish (*Danio rerio*) adults and embryos were maintained in accordance with the Animals (Scientific Procedures) Act 1986 under license from the United Kingdom Home Office (PPL7266180). Wildtype AB strain zebrafish were maintained in the Fish Facility at King’s College London in a recirculating system containing fish water with a zinc concentration of 0.060 ± 0.004 μM, at 28.5 °C under a 14 h light/10 h dark photoperiod. Embryos were obtained by placing sexually mature males and females in breeding tanks the evening prior to collection. Fertilised embryos were collected the following morning and transferred into 1× Danieau solution (58 mM NaCl, 0.7 mM KCl, 0.4 mM MgSO_4_, 0.6 mM Ca(NO_3_)_2_, 5 mM HEPES, pH 7.6). For experiments involving larvae younger than 5 days post-fertilisation (dpf), embryos were maintained at 28.5 °C in Petri dishes at a density of approximately 60 embryos per dish.

### Generation of *zip9* knockout zebrafish

A CRISPR/Cas9 genome-editing strategy was used to generate a *zip9* knockout zebrafish line. Guide RNA sequences targeting exon 1 of the *zip9* gene were designed using CHOPCHOP and are listed in Supplementary Table S1. A target site located in exon 1 of the *zip9* gene was selected (Supplementary information Table 1). The crRNA and tracrRNA were annealed to generate a gRNA duplex and subsequently incubated with Cas9 protein to form a ribonucleoprotein (RNP) complex. Approximately 2 nL of RNP mixture was injected into the cytoplasm of one-cell-stage embryos within 30 min post-fertilisation. Genomic DNA was extracted from larvae or fin clips by alkaline lysis followed by neutralisation. PCR amplification and Sanger sequencing were used to assess editing efficiency and identify mutant alleles. A heterozygous founder (F0) was out-crossed with wildtype fish to generate F1 offspring, which were subsequently used to establish the mutant line.

### RNA isolation and quantitative real-time PCR

Total RNA was extracted using TRIzol reagent (Invitrogen) according to the manufacturer’s instructions. RNA concentration and purity were determined using a NanoDrop spectrophotometer. Reverse transcription was performed using 500 ng RNA with the High-Capacity cDNA Reverse Transcription Kit (Thermo Fisher Scientific). Quantitative real-time PCR (qPCR) reactions were performed using Luna Universal qPCR Master Mix (New England Biolabs) on a QuantStudio 7 Real-Time PCR system. Each reaction contained 5 µL master mix, 0.5 µL of each primer (10 µM), and diluted cDNA in a total volume of 10 µL. Cycling conditions consisted of 95 °C for 1 min, followed by 40 cycles of 95 °C for 15 s, 60 °C for 30 s, and 72 °C for 30 s, with a final melting curve analysis. Primer efficiencies were validated using standard curves generated from serial dilutions. Primer sequences are listed in Supplementary information Table 2.

### Western blot analysis

Protein extracts (30 µg per sample) were separated by SDS-PAGE and transferred onto nitrocellulose membranes. Membranes were incubated with primary antibodies against Zip9 (kind gift from Prof. Peter Thomas, previously described^**24**^) and GAPDH (Sigma-Aldrich), followed by HRP-conjugated secondary antibodies. Protein bands were visualised using an enhanced chemiluminescence (ECL) detection system.

### Whole-mount immunofluorescence staining

To determine Zip9 localisation in zebrafish larvae, whole-mount immunofluorescence staining was performed at 5 dpf. Larvae were euthanised using MS-222 and fixed in 4% paraformaldehyde (PFA). Samples were bleached, permeabilised, and blocked with 5% normal goat serum, followed by incubation with anti-Zip9 primary antibody (1:400) and Alexa Fluor-conjugated secondary antibody (1:500). Larvae were mounted in low-melting agarose and imaged using confocal microscopy.

### LA-ICP-MS analysis of larval metal distribution

Elemental distribution in zebrafish larvae was analysed using laser ablation inductively coupled plasma time-of-flight mass spectrometry (LA-ICP-ToF-MS). 5 dpf zebrafish larvae were euthanised using tricaine methane sulfonate (MS-222) and embedded in optimal cutting temperature compound (OCT), snap-frozen in powdered dry ice, and cryo-sectioned at 15 µm thickness. Elemental imaging was performed at the London Metallomics Facility using an Iridia 193 nm ArF excimer laser ablation system (Teledyne Photon Machines, USA) coupled via an Aerosol Rapid Introduction System (ARIS) to a Vitesse ICP-TOF-MS (Nu Instruments, UK). Imaging was conducted in fixed dosage mode (15) with a spatial resolution of 10 µm and a repetition rate of 1000 Hz. Seventeen elements were measured simultaneously, including Na, Mg, Al, Si, P, S, Cl, K, Ca, V, Cr, Mn, Fe, Co, Ni, Cu, and Zn. Elemental images were reconstructed by combining ICP-MS signal intensities with spatial coordinates using HDF-based Image Processing software. Representative elemental distribution maps for all measured elements in wildtype and *zip9* mutant larvae are provided in Supplementary Fig. S2 and S3 to illustrate the overall spatial distribution of metals detected by LA-ICP-MS.

### Synchrotron X-ray fluorescence imaging

Zebrafish larvae at 5 days post fertilisation (5 dpf) were euthanised using MS-222. Approximately 40% of the posterior tail region was removed for genomic DNA extraction and genotyping by Sanger sequencing. The remaining portion of each larva, containing the head and anterior body, was embedded in OCT compound, mounted in 0.55 mm polyimide capillaries (Goodfellow), snap frozen, and stored at −80 °C prior to analysis. Elemental imaging was performed using synchrotron-based X-ray fluorescence (XRF) at beamline I18 of Diamond Light Source^**34**^ (Didcot, UK). Elemental maps were acquired using an incident beam energy of 11 keV focused to a ~2 μm2 spot on the sample using Kirkpatrick-Baez mirrors. Two Vortex®-ME4 Silicon Drift Detectors (Hitachi High-Tech, Japan) and the Xspress3 X-ray fluorescence detector readout system (Quantum Detectors, UK) were used with a dwell time of 0.5 s per pixel. For two-dimensional imaging, samples were scanned at two orientations corresponding to dorsal and lateral views. The 0° orientation (dorsal view) was defined as the position in which both eyes and otoliths were symmetrically visible in the projection image, indicating that the dorsal side of the head was facing the detector. The 90° orientation (lateral view) was obtained by rotating the sample by approximately 90° relative to this reference position, resulting in the two eyes appearing overlapped in projection and allowing visualisation of the lateral side of the head. For one wildtype sample, a dwell time of 0.1 s per pixel was used for the 90° scan. 2D elemental maps were analysed using Fiji (ImageJ). Zinc and calcium channels were displayed using an intensity range of 0-4000 arbitrary units (a.u.), pseudo-coloured (Zn, green; Ca, red), and merged for visualisation. Regions of interest (ROIs) were defined using the eyes and otoliths as anatomical landmarks. The ROI extended from the dorsal edge of the eyes ventrally by approximately two eye diameters and spanned the width between the lateral edges of both eyes, while excluding the eye regions to focus analysis on the hypothalamus-pituitary region. Mean grey values within ROIs were extracted to represent elemental signal intensity and were normalised to dwell time (a.u./s) to allow comparison between scans acquired with different dwell times. For three-dimensional XRF tomography a single slice was measured by translating the sample horizontally and rotating 180° around the vertical axis. The sample was then translated vertically 15 µm and the measurement repeated, when stacked together these slices provided volumetric information. Image reconstructions were performed using the iterative technique maximum-likelihood expectation maximization algorithm^**46**^. Full sample data sets were rendered using Avizo software (version 2021.1, Thermo Fisher Scientific).

### Reproductive assays and embryo development analysis

To assess reproductive performance, sexually mature zebrafish (3.5 months old) were paired in breeding tanks. Males and females were separated overnight and allowed to spawn after barrier removal the following morning. Spawning behaviour was monitored, and the spawning rate and egg numbers were recorded. Embryonic development was monitored following fertilisation. Embryos were incubated in Danieau solution and observed at defined developmental time points up to 24 hours post-fertilisation (hpf) to assess developmental progression and survival.

### Gonad histology

Gonads from wildtype and *zip9* mutant zebrafish were fixed and dissected in 4% PFA. Samples were dehydrated through graded ethanol, cleared in Histo-Clear, and embedded in paraffin wax. Sections (5 µm) were prepared using a microtome and stained with haematoxylin and eosin (H&E) for histological analysis.

### Statistical analysis

All statistical analyses were performed using GraphPad Prism 10. Data were analysed using unpaired t-tests, one-way ANOVA, or two-way ANOVA followed by appropriate multiple comparison tests (Dunnett’s, Tukey’s, or Šidák’s). Data are presented as mean ± SEM, and statistical significance was defined as *P* < 0.05. All experiments were performed with at least three independent biological replicates unless otherwise stated.

## Acknowledgements

We thank the staff of the King’s College London Fish Facility for zebrafish husbandry and animal care. We acknowledge Diamond Light Source for access to beamline I18 (proposal number SP39141), which contributed to the results presented here. We also acknowledge the London Metallomics Facility for support with LA-ICP-MS analysis. We are grateful to Peter Thomas for providing the Zip9 antibody used in this study.

