## Supplementary information for "Deficiency of the membrane androgen receptor ZIP9 alters brain zinc distribution, reproductive endocrinology, and female fertility"

**Supplementary Table S1**

CRISPR/Cas9 guide RNA and genotyping primers used for generation of the *zip9* mutant zebrafish line.

The table lists the guide RNA sequence targeting exon 1 of the zebrafish *zip9* gene and the forward and reverse primers used for mutation screening and genotyping.

| gRNA | TCAGAGAAGTTTACAGCCAA |
| --- | --- |
| Forward primer for screening | CCATCAGTCTGCTGTCTCTGTC |
| Reverse primer for screening | GCATAGTCCCAAGGAGATTCAG |

**Supplementary Table S2**

Primer sequences used for quantitative real-time PCR analysis.

Forward (Fwd) and reverse (Rev) primer sequences used for qPCR analysis of zebrafish genes examined in this study.

| Gene | Primer sequence (5’_3’) | GeneBank accession |
| --- | --- | --- |
| *18s* | Fwd: TCGCTAGTTGGCATCGTTTATG  Rev: CGGAGGTTCGAAGACGATCA | XR_012407107.1 |
| *zip9* | Fwd: CATCTATTAGTCTTTGCTCTGG  Rev: CTCTTTACTGCTCTGACTGA | NM_001013540.1 |
| *ar* | Fwd: ACAACACACCTGGATGGGAGTGAT  Rev: TGACCTGTAGCAGCACAAACTCCT | NM_001083123.1 |
| *kiss1* | Fwd: TCTAAACTCTCAGCGCTCTTCT  Rev: TGTCCTGTTCTCTCTTGCCATA | NM_001113489.1 |
| *kiss2* | Fwd: CGACTCTGACAGACTCAAACAC  Rev: AGAAAATCGCATCCTTCTGACG | NM_001142585.1 |
| *gnrh2* | Fwd: CAGAGGTTTCAGAGGAAGTGAAGC  Rev: TGAGGGCATCCAGCAGTATTG | NM_181439.4 |
| *gnrh3* | Fwd: CACAACAGCAACAAAGGTGATTC  Rev: CCAGATGCCCAGCAGGTAAT | NM_001177450.1 |
| *fshb* | Fwd: TGATTCAGTCTTCGTGTACCCC  Rev: AGTGCTCTAGTGTATGCTGCAG | NM_205624.2 |
| *lhb* | Fwd: GTGCACCATAAACACTTCCGAC  Rev: CTCGACTGTGTGTGTAGGTTGA | NM_205622.2 |
| *esr1* | Fwd: ACTCTCACCCATGTACCCTAAGG  Rev: CGGGTAGTATCCCACTGAAGC | NM_152959.1 |
| *fshr* | Fwd: AACATGCACATAGAGAGGATTCCCAG  Rev: GCTCAGTAAACAGCTCCAGGC | NM_001001812.1 |
| *lhr* | Fwd: ATCACTCACGCTCTCCGACT  Rev: GCTGCTGACGCCTATTAAGG | NM_001001812.1 |
| *cyp19a1* | Fwd: GTTCAGTCTTGACAACTTCCATAAAAAT  Rev: TGCGACAGGTTGTTGGTTTGC | NM_131154.4 |
| *yap1* | Fwd: ACGGGTGGGAACAAGCTATT  Rev: CCTTGCTTTACTGGGGCACT | NM_001139480.1 |
| *vtg* | Fwd: CCTTGGAGAAAATTGAGGCTATC  Rev: CTGAATGAACTCGGGAGTGGTA | NM_001044897.3 |

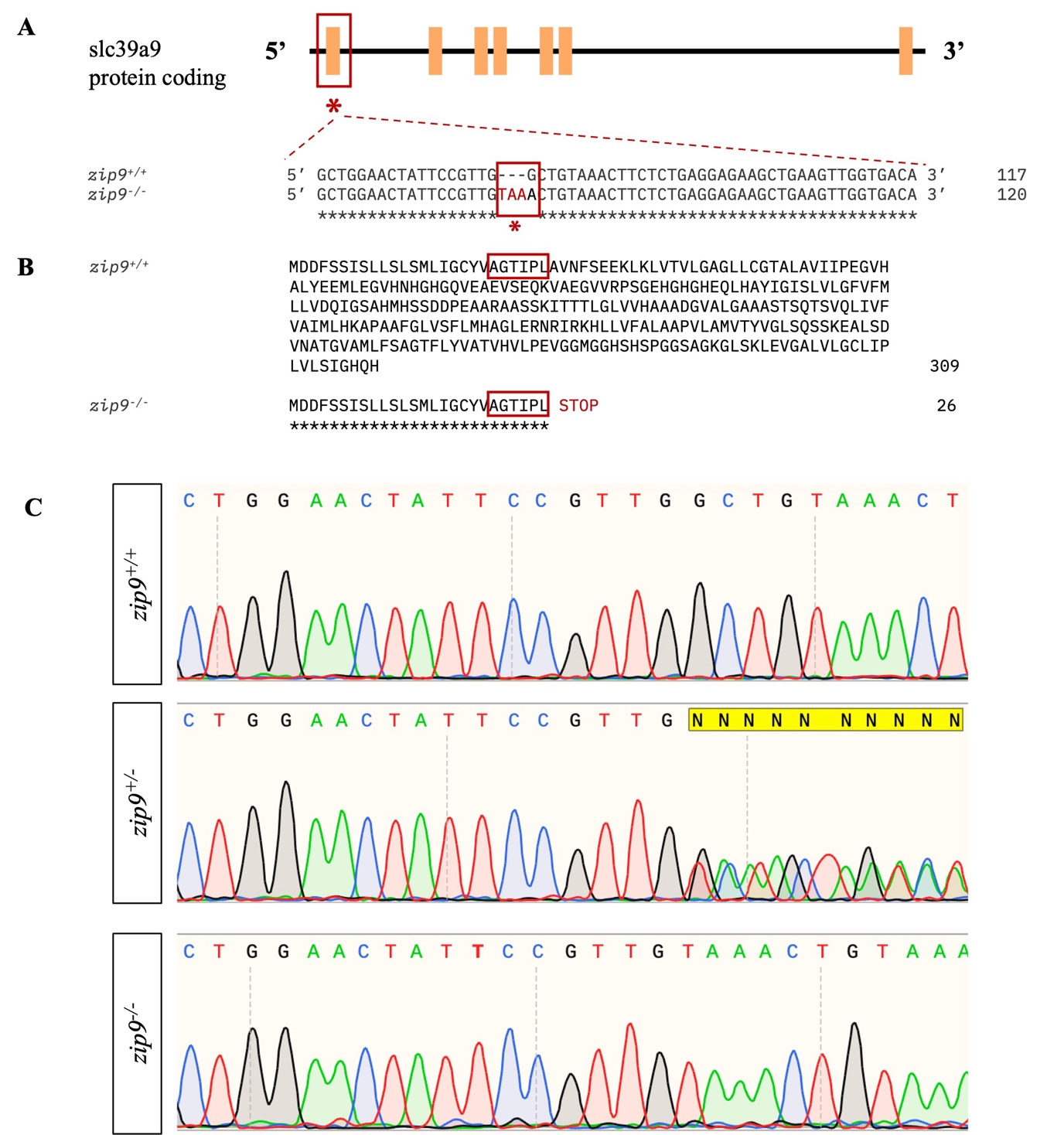

**Supplementary Figure S1.** Generation of the *zip9* knockout zebrafish line. (A) Schematic representation of the *zip9* gene showing the CRISPR/Cas9 target site in exon 1. A four-base-pair insertion introduced a frameshift mutation and premature stop codon (TAA). (B) Predicted truncation of the Zip9 protein following the mutation. (C) Representative Sanger sequencing chromatograms showing wildtype (*zip9^+/+^*), heterozygous (*zip9^+/-^*), and homozygous mutant (*zip9^-/-^*) genotypes.

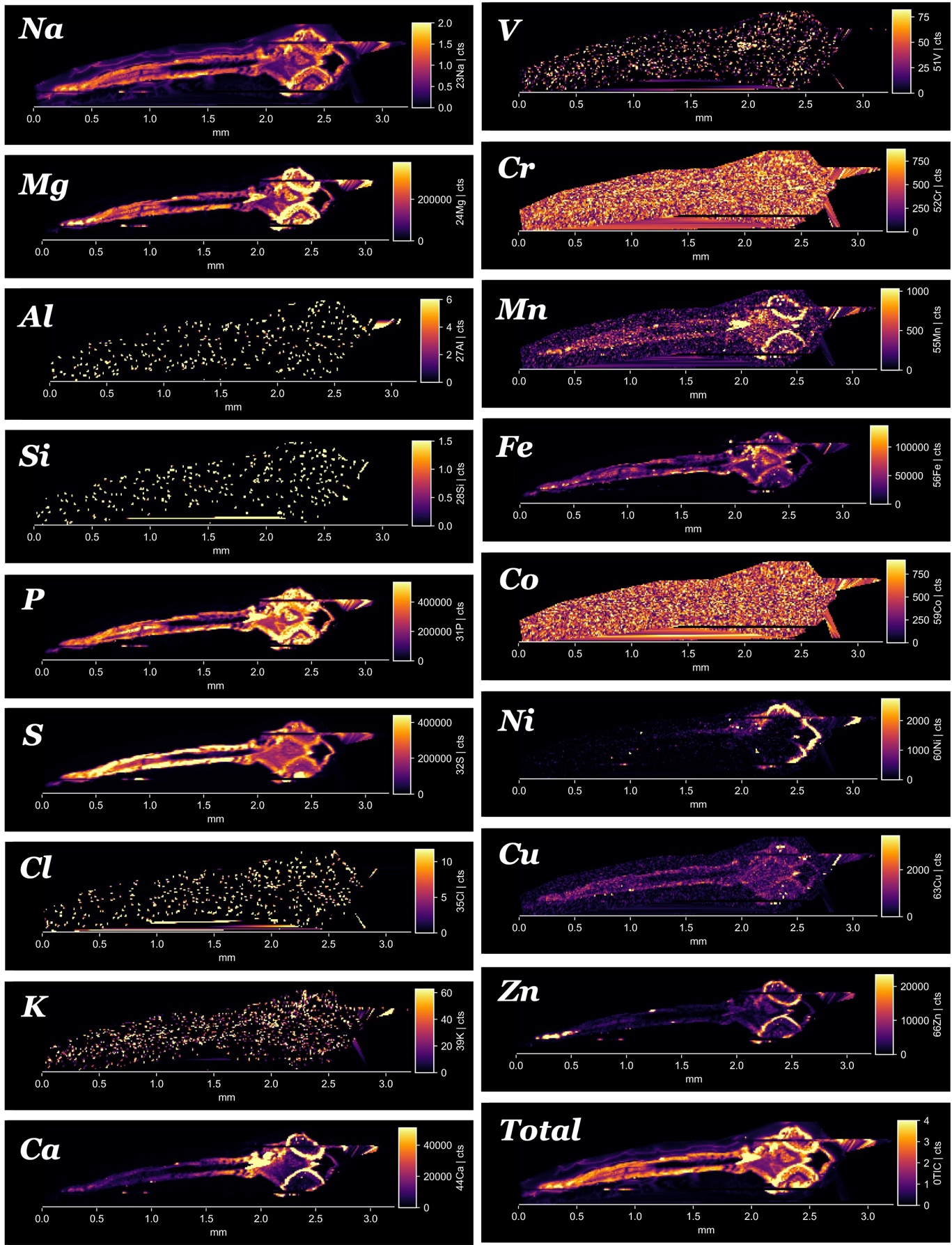

**Supplementary Figure S2.** Representative LA-ICP-MS elemental maps of wildtype zebrafish larvae. Elemental distributions of sodium (Na), magnesium (Mg), aluminium (Al), silicon (Si), phosphorus (P), sulphur (S), chlorine (Cl), potassium (K), calcium (Ca), vanadium (V), chromium (Cr), manganese (Mn), iron (Fe), cobalt (Co), nickel (Ni), copper (Cu), and zinc (Zn) in 5 dpf wildtype zebrafish larvae. Elemental images are displayed as relative signal intensity.

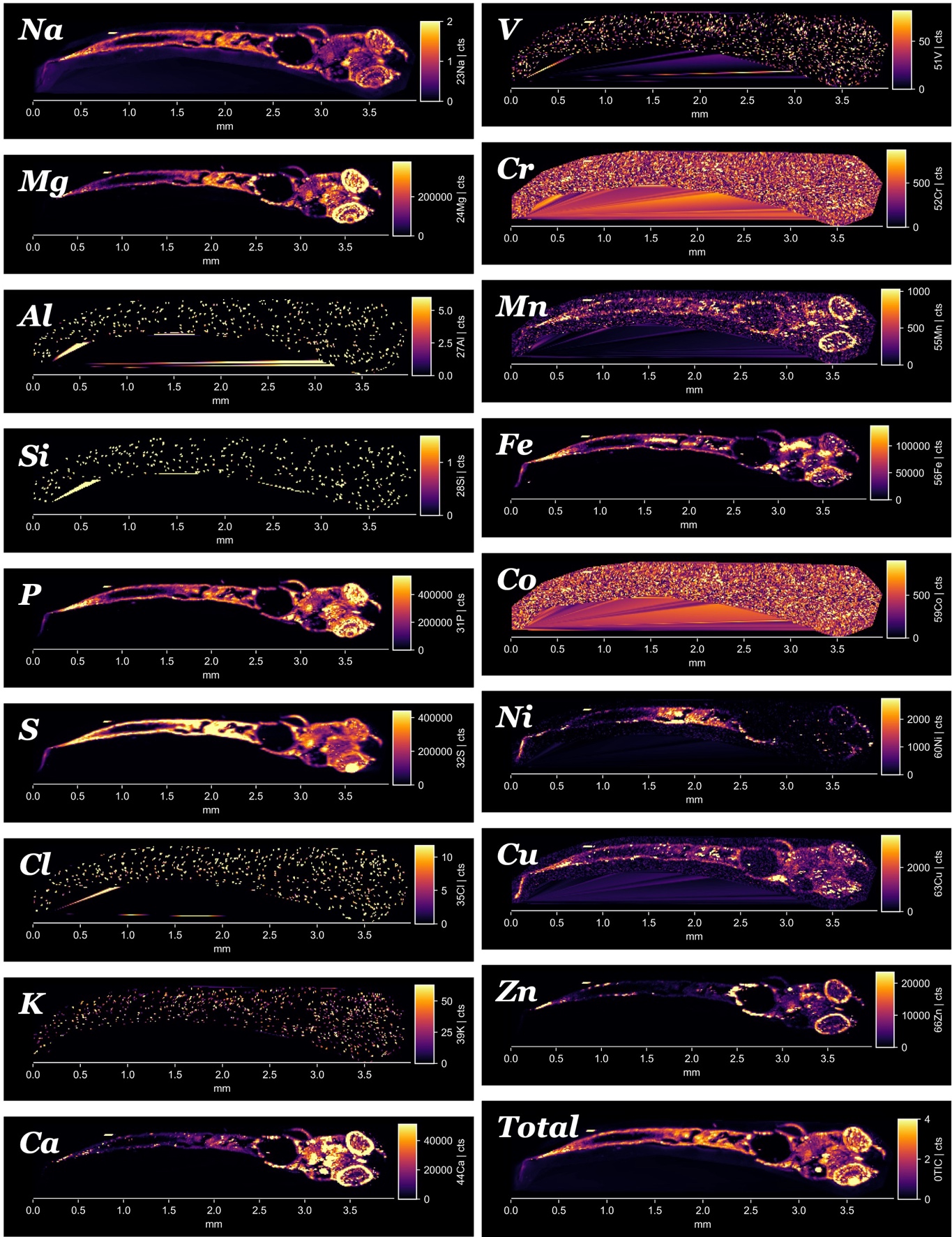
**Supplementary Figure S3.** Representative LA-ICP-MS elemental maps of *zip9^-/-^* zebrafish larvae. Elemental distributions of sodium (Na), magnesium (Mg), aluminium (Al), silicon (Si), phosphorus (P), sulphur (S), chlorine (Cl), potassium (K), calcium (Ca), vanadium (V), chromium (Cr), manganese (Mn), iron (Fe), cobalt (Co), nickel (Ni), copper (Cu), and zinc (Zn) in 5 dpf *zip9^-/-^* zebrafish larvae. Elemental images are displayed as relative signal intensity.

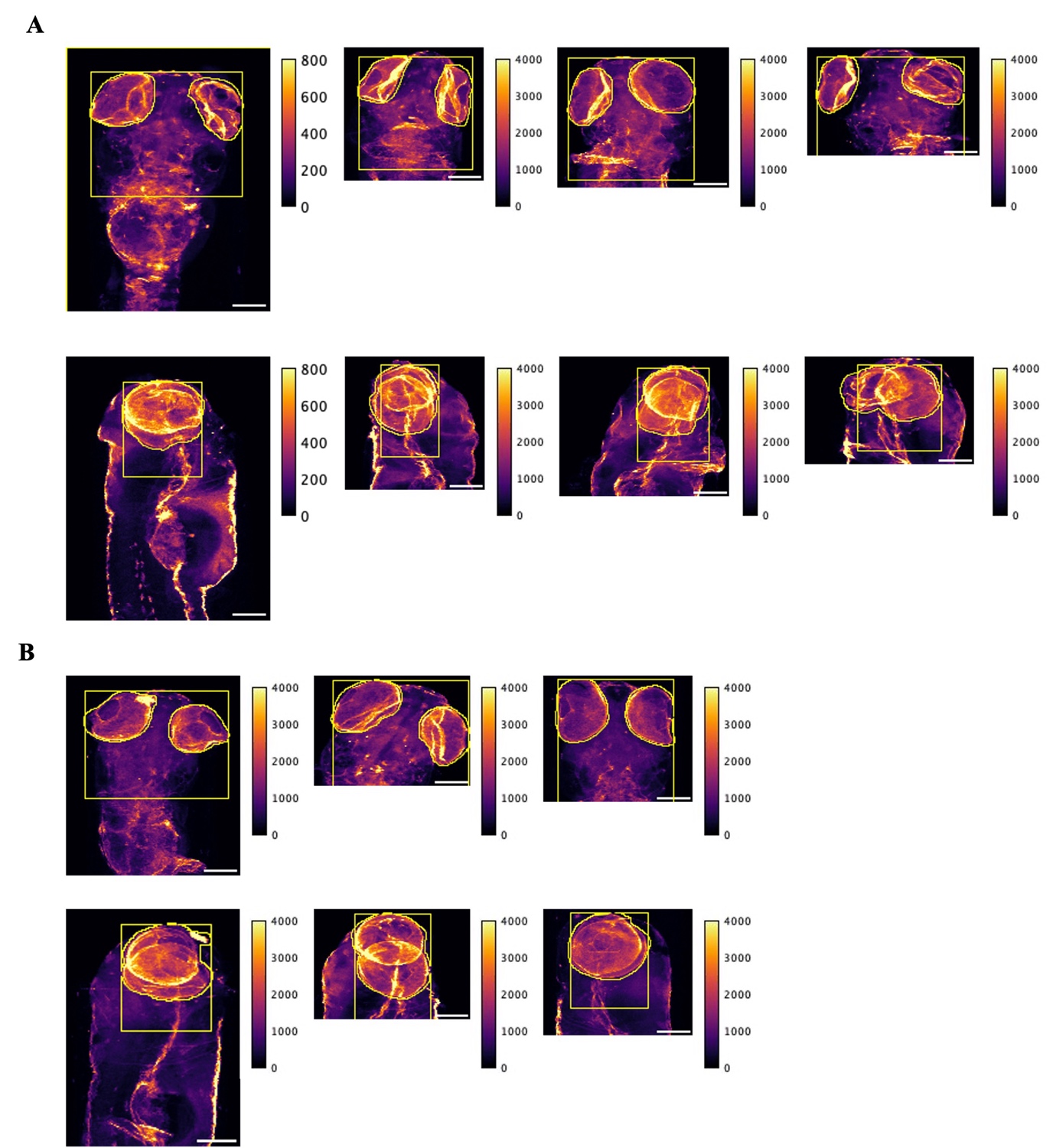

**Supplementary Figure S4.** Representative X-ray fluorescence (XRF) maps of zebrafish heads used for ROI analysis. Two-dimensional XRF scans showing zinc and calcium distribution in wildtype and *zip9* mutant zebrafish heads. Samples were scanned at 0° and 90° orientations. The ROI used for quantification of elemental intensity in the hypothalamus-pituitary region are indicated. Multiple biological samples are shown for each genotype. Scale bar: 0.2 mm.

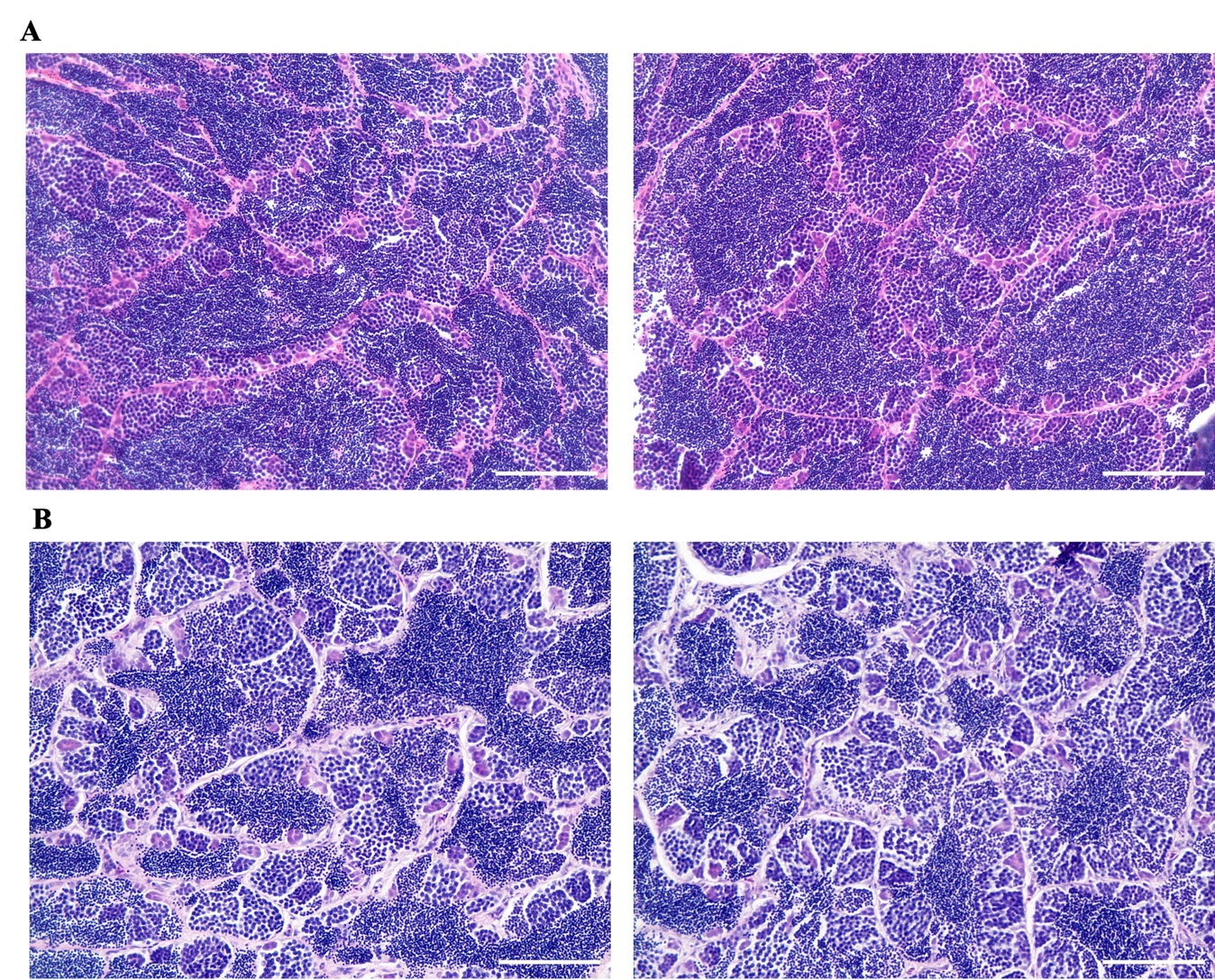

**Supplementary Figure S5.** Histological analysis of testes from adult zebrafish. Representative haematoxylin and eosin (H&E) stained sections of testes from wildtype (A) and *zip9* mutant (B) zebrafish. Scale bar: 100 µm.
